# Multi-objective computational optimization of human 5′ UTR sequences

**DOI:** 10.1101/2024.11.12.622374

**Authors:** Keisuke Yamada, Kanta Suga, Naoko Abe, Koji Hashimoto, Susumu Tsutsumi, Masahito Inagaki, Fumitaka Hashiya, Hiroshi Abe, Michiaki Hamada

**Affiliations:** Department of Electrical Engineering and Bioscience, Graduate School of Advanced Science and Engineering, Waseda University, Tokyo 169-8555, Japan; Department of Bioengineering, University of Pennsylvania, Philadelphia, PA 19104, USA; Computational Bio Big-Data Open Innovation Laboratory, AIST-Waseda University, Tokyo 169-8555, Japan; Graduate School of Medicine, Nippon Medical School, Tokyo 113-8602, Japan; Department of Chemistry, Graduate School of Science, Nagoya University, Furo-cho, Chikusa-ku, Nagoya, Aichi 464-8602, Japan; Graduate School of Arts and Sciences, The University of Tokyo, 3-8-1 Komaba, Meguro-ku, Tokyo 153-8902, Japan; Research Center for Materials Science, Nagoya University, Furo-cho, Chikusa-ku, Nagoya, Aichi 464-8602, Japan; Institute for Glyco-core Research (iGCORE), Nagoya University, Furo-cho, Chikusa-ku, Nagoya, Aichi 464-8601, Japan

## Abstract

The computational design of mRNA sequences is a critical technology for both scientific research and industrial applications. Recent advances in prediction and optimization models have enabled the automatic scoring and optimization of 5′ UTR sequences, key upstream elements of mRNA. However, fully automated design of 5′ UTR sequences with more than two objective scores has not yet been explored. In this study, we present a computational pipeline that optimizes human 5′ UTR sequences in a multi-objective framework, addressing up to four distinct and conflicting objectives. Our work represents an important advancement in the multi-objective computational design of mRNA sequences, paving the way for more sophisticated mRNA engineering.

## 1 Introduction

The success of messenger RNA (mRNA) vaccines against SARS-CoV-2 (Polack *et al*., 2020; Baden *et al*., 2021) has significantly heightened interest in mRNA design within both the scientific community and industry. Among various design parameters, the mRNA sequence itself offers a vast combinatorial design space, expanding exponentially with increasing length. Several studies have presented rational mRNA design strategies based on thermodynamic energy scoring (Mauger *et al*., 2019; Wayment-Steele *et al*., 2021). The growing attention to mRNA design also extends to specific mRNA components, including the 5′ cap (Inagaki *et al*., 2023) and the sequence composition of untranslated regions (UTRs).

The 5′ UTR, an upstream element of mRNA, plays essential biological roles such as regulating translation (Sonenberg and Hinnebusch, 2009; Hinnebusch *et al*., 2016) and stabilizing mRNA (Jia *et al*., 2020). Recognizing the importance of UTRs for mRNA-based therapeutics, Asrani *et al*. (2018) and Leppek *et al*. (2022) have explored combinations of known 5′ UTRs and 3′ UTRs to enhance protein expression and mRNA stability. These studies highlight that designing 5′ UTRs with desired characteristics is a key step in optimizing mRNA design.

Recent studies have explored fully computational approaches for designing 5′ UTR sequences, leveraging largescale empirical datasets and the deep learning model named Optimus 5-Prime, introduced by Sample *et al*. (2019). The genetic algorithm, also presented by Sample *et al*. (2019), laid the groundwork for model-based optimization of 5′ UTR sequences, using Optimus 5-Prime as the primary oracle. Subsequent methods, including the deep exploration network (Linder *et al*., 2020) and gradient-based activation maximization (Linder and Seelig, 2021; Castillo-Hair and Seelig, 2022), furthered the exploration of 5′ UTR sequence. Additionally, researchers in computer science have developed model-based optimization techniques, such as the cross-entropy method (Gane *et al*., 2019), conservative objective model (Trabucco *et al*., 2021), and iterative posterior scoring (Zhang *et al*., 2022). These studies highlight the growing diversity of computational 5′ UTR sequence design methods; however, most focus on a single optimization target to improve mRNA trans-lation levels.

The real-world challenges of mRNA sequence design involve multiple requirements, making single-target optimization insufficient. The LinearDesign algorithm introduced by Zhang *et al*. (2023) was an early effort to balance both mRNA stability and codon optimality. While relatively few studies have addressed mRNA sequence design, multi-objective sequence optimization has been being an attractive topic in computational protein sequence design (Stanton *et al*., 2022; Jain *et al*., 2023; Gruver *et al*., 2023). Therefore, the multi-objective optimization of 5′ UTR sequences remains a crucial yet achievable challenge.

Here, we present a multi-objective computational design of human 5′ UTR sequences. By leveraging latent representations from a pre-trained DNA language model and a latent-based multi-objective Bayesian optimization, we optimized human 5′ UTR sequences across four design objectives. We also compared different candidate selection methods, an important step in practice for selecting candidates for experimental testing. Our study offers a comprehensive workflow of human 5′ UTR sequence design from defining a multi-objective optimization problem to experimental validation.

## 2 Materials and methods

### 2.1 LaMBO models

We employed latent-based multi-objective Bayesian optimization (LaMBO) (Stanton *et al*., 2022) to optimize sequences with multiple objectives. The foundational LaMBO model integrates a convolutional neural network (CNN)-based autoencoder with a Gaussian process (GP) head. In addition to the original architecture, two additional models were incorporated. The CNN-based autoencoder was replaced with a bidirectional encoder representation from transformers (BERT) (Devlin *et al*., 2018), either with randomly initialized parameters or with those from DNABERT (Ji *et al*., 2021), a DNA language model pre-trained on the human genome. To leverage the pre-trained parameters, the sequence representation generated by the DNABERT encoder was directly passed to the DNABERT decoder, omitting the concatenation to the processed representation for the GP head. Throughout this paper, each model is referred to by its autoencoder architecture, such as LaMBO-CNN, LaMBO-BERT, and LaMBO-DNABERT. The architecture and training procedure of LaMBO-BERT and LaMBO-DNABERT are summarized in Fig. 1. For detailed mathematical descriptions, we refer readers to the original work by Stanton *et al*. (2022).

**Figure 1:**
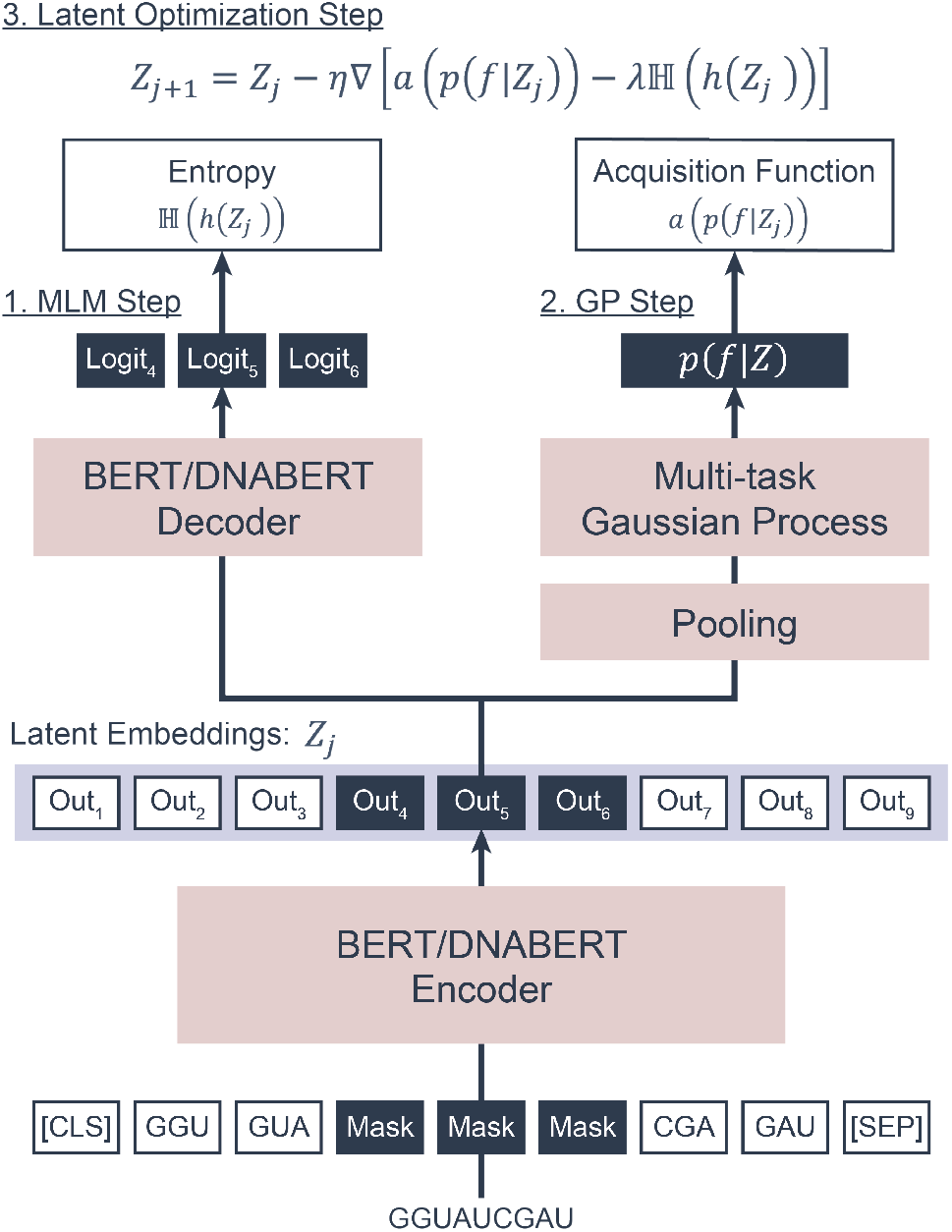
Our multi-objective optimization process sequentially updates the pool of 5′ UTR sequences across multiple rounds, with each round comprising three steps. First, the masked language modeling (MLM) step trains the sequence encoder and decoder using a MLM objective to generate latent embeddings and enhance the model’s decoding capacity. Pretrained parameters from DNABERT are introduced in this step to aid the training process. Second, the multi-task Gaussian process head is trained using independently predicted objective values, including AGC content, mean ribosome load, G4 score, and *in vitro* stability. Once trained, the parameters shown in red in the figure are frozen, and the latent embeddings *Zj* are directly optimized. Third, the latent optimization step aims to increase the likelihood of score improvement, measured by the acquisition function *a*(·), while avoinding a uniform proposal distribution by applying an entropy penalty *λ*ℍ(·), where *λ* is the penalty parameter, and ℍ(·) is the Shannon entropy. In the figure, *h*(·) denotes the decoder, *p*(*f*|*Z*) represents the posterior predictive distribution, and *η* represents the step size of latent optimization.

### 2.2 Sequence design objectives

We aimed to design 5′ UTR sequences of human mRNAs using a fully computational approach, emphasizing multi-objective criteria. Our 5′ UTR design target, established by Sample *et al*. (2019), focused on the 50 nucleotides upstream of the start codon. To utilize available prediction models for 5′ UTR properties, we focused on designing sequences with unmodified bases as discussed below. Following their design, we set four design objectives as follows.

#### 2.2.1 Mean ribosome load (MRL)

The mean ribosome load (MRL) indicates the average number of ribosomes on an mRNA sequence. In earlier work, Sample *et al*. (2019) generated sequence-MRL datasets from randomized 5′ UTRs and trained a CNN model, named Optimus 5-Prime, to predict MRL based on the 5′ UTR sequence. Since ribosome occupancy of mRNA correlates with protein levels (Cenik *et al*., 2015), MRL could serve as a proximate metric for the translation level of the mRNA. Consequently, we adopted Optimus 5-Prime as an objective function to maximize.

#### 2.2.2 AGC content

*In vivo* delivered mRNAs, particularly those with high uridine content, tend to stimulate innate immune response in human cells (Karikó *et al*., 2004; Kawai and Akira, 2010). One approach to diminish mRNA immunogenicity is through chemical modifications such as replacing uridines with pseudouridines (Karikó *et al*., 2008). However, the effect of such chemical modifications on the translation level of the delivered mRNAs remains debatable (Uchida *et al*., 2015; Thess *et al*., 2015; Kauffman *et al*., 2016). Some studies suggest that uridine depletion is more effective in augmenting the activity of translated proteins than chemical modifications of mRNAs (Vaidyanathan *et al*., 2018). In light of these findings, one of our objectives was to reduce the uridine content of 5′ UTR, aiming to curb the immunogenicity of mRNAs from the nucleotide sequence design perspective.

#### 2.2.3 *in vitro* stability

Stabilizing mRNA is crucial for both the efficient translation of the target gene (Jia *et al*., 2020) and the efficient production and storage of mRNA-based therapeutics (Zhao *et al*., 2020). Wayment-Steele *et al*. (2022) organized a Kaggle competition centered on predicting *in vitro* mRNA degradation at the nucleotide level. We employed the winning model, Nullrecurrent, and used the average value over the mRNA sequence as our stability measure to optimize.

#### 2.2.4 G-quadruplex

The G-quadruplex (G4) is a rigid structure found in DNA and RNA sequences rich in guanines. Although its precise biological roles remain unclear, G4s in 5′ UTRs have been linked to translational inhibition (Varshney *et al*., 2020; Beaudoin and Perreault, 2010; Murat *et al*., 2018). Since our objectives tend to elevate the GC content both directly and indirectly, the resulting sequence is prone to form G4s. To counter this, we incorporated a G4 score predicted by the deep learning prediction model, DeepG4 (Rocher *et al*., 2021) and set an objective to minimize the G4 score. The optimization of G4 score, however, did not require the same rigor as our other objectives, so we relaxed its optimization. Specifically, the DeepG4 output, initially a real value between 0 and 1, was quantized at intervals of 0.1.

### 2.3 Sequence optimization

#### 2.3.1 Dataset

The dataset from Sample *et al*. (2019)’s study, which was measured using the eGFP gene with unmodified uridines (Accession number: GSM3130435), underwent the following preprocessing. First, sequences with zero sequencing reads in any polysome fractions were excluded. The top 512 sequences based on the highest total read counts, were chosen to form the initial sequence pool (pool 1) for hyperparameter tuning. Subsequently, three additional sets of 512 sequences with highest total read counts were used as initial sequence pools (pools 2-4) during model evaluation and sequence optimization.

#### 2.3.2 Model training

For each model, a selected set of hyperparameters (Supplementary Table S1) was tuned using the initial sequence pool 1 with the grid search method to optimize the relative hypervolume averaged over five seeds. In addition, LaMBO-DNABERT were trained while linearly decreasing the number of masked tokens from a defined mask ratio to one token per sequence, promoting exploration in earlier rounds and gradually enhance exploitation over time. LaMBO-BERT was also trained in the same manner as LaMBO-DNBERT. Models with the tuned hyperparameters were then trained using the distinct three initial sequence pools (pools 2-4) for 64 rounds.

#### 2.3.3 Candidate selection

To select candidate sequences for experimental validation, we used a classical Pareto ranking method (Carlos M. Fonseca, 1993). Each individual’s rank score was calculated by the number of solutions in the population that the individual dominates, and those with the highest rank scores were selected. To ensure the diversity of selected sequences, optimized sequences from the three different initial pools (pools 2-4) were combined together. The sequences were then clustered using UCLUST (Edgar, 2010) with a sequence identity threshold of 60%, and only one sequence with the highest Pareto ranking score per cluster was selected.

### 2.4 Experimental evaluation

Each selected 5′ sequence was concatenated to the 5′ end of an eGFP coding sequence and a poly-A sequence, then amplified with an upstream T7 promoter by PCR to prepare template DNA for *in vitro* transcription (IVT). From the template DNA, 5′ capped mRNAs were prepared by IVT using a hydrophobic-tagged dinucleotide cap analog, DiPure (Inagaki *et al*., 2023). The mRNA was mixed with Lipofectamine MessengerMAX reagent (Thermo) in Opti-MEM I Reduced Serum Media (Thermo) and transfected into 293T cells (Riken Cell Bank, RCB2202) in a 96-well plate. Fluorescence (excitation at (485 ± 10) nm, emission at (535 ± 10) nm) was measured at 30-minute intervals for 24 hours after transfection. Measured values were summed to determine the expression level of the mRNA. Detailed experimental descriptions are provided in the supplementary methods.

## 3 Results

To apply multi-objective Bayesian optimization to 5′ UTR sequences, we initially employed LaMBO-CNN with two objectives; increasing MRL and AGC content. The Pareto frontier, in terms of MRL and AGC content, continuously expanded over 64 optimization rounds (Fig. 2a). When evaluated for additional two metrics, G4 score and *in vitro* stability, the non-dominated solutions from the LaMBO-CNN with two objectives exhibited suboptimal values, even in the initial sequence pool. On the other hand, when these additional objectives were integrated into the model, LaMBO-CNN was able to balance all four objectives while maintaining some solutions with comparable performance for the original two objectives (Fig. 2b). These results demonstrated that 5′ UTR sequences can be computationally optimized with up to four objectives in parallel.

**Figure 2:**
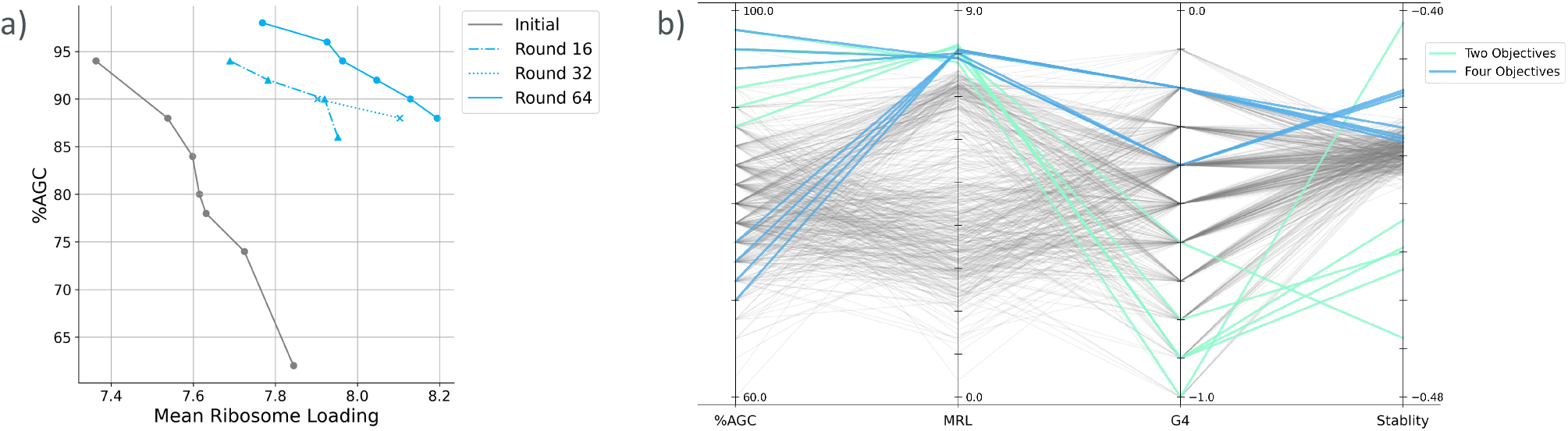
a) LaMBO-CNN was trained over 64 rounds with AGC content and predicted mean ribosome load (MRL) as the objectives. The Pareto frontier from each round is visualized as a connected line. b) Comparison of two-objective and four-objective optimizations. Colored lines indicate the non-dominated solutions, in terms of AGC content and MRL, obtained after 64 rounds of training LaMBO-CNN with two and four objectives. The non-dominated solutions shown in a) are further labeled with predicted G4 scores and *in vitro* stability. Gray lines represent the scores of all 512 sequences in the initial sequence pool.

Building upon the original architecture, we tested LaMBO with different encoders. We used a BERT encoder with random initial parameters and a DNABERT encoder, pre-trained on the human genome, intending to leverage both the larger parameter size and the genomic knowledge acquired during pre-training. After hyperparameter tuning on a set of sequences, each model was evaluated by optimizing three distinct sets of sequences (Methods 2.3.1). Among the three models, LaMBO-DNABERT showed the highest average performance, as measured by the relative hypervolume of the four objective scores (Fig. 3). We also analyzed the model’s decoding capacity for masked tokens by measuring normalized perplexity. Compared to models with randomized initial parameters, LaMBO-DNABERT maintained low perplexity, indicating it effectively utilized the original decoding capacity of DNABERT (Supplementary Figure S1).

**Figure 3:**
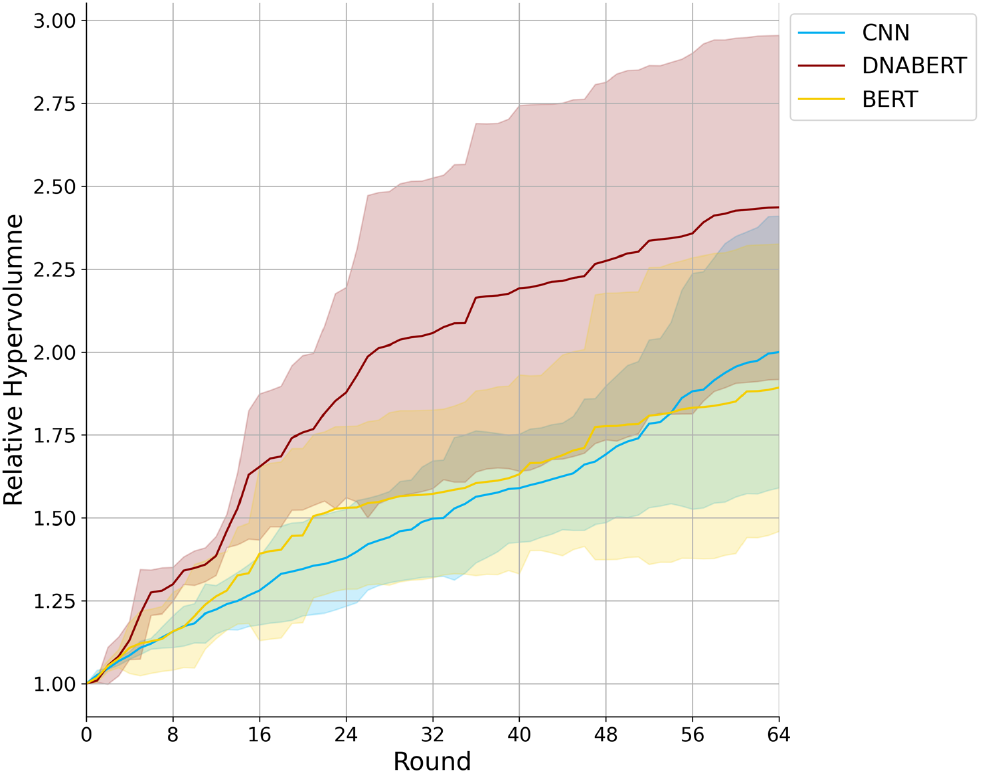
Each model was trained to optimize three sets of initial sequences over 64 rounds, with performance measured by the relative hypervolume across four objectives. The shaded areas represent the confidence intervals for the three sets. Note that the hyperparameters of each model were tuned using a separate set of initial sequences.

After sequence optimization, only a small subset of candidates can be evaluated through labor-intensive experiments, making selection a non-trivial problem in multi-objective optimization. As the number of objectives increases, the number of non-dominated solutions grows, complicating the ranking process. Ranking non-dominated solutions requires assigning weights or ranks to objectives, often involving subjective decisions. To address these challenges, we relied on the Pareto ranking method (Carlos M. Fonseca, 1993). The Pareto ranking method treats every objective equally, imposes no quantitative assumptions among objectives, and is subjective to the score distribution within the population. We reasoned that it would be well-suited to our purpose, given that we were dealing with sequences already explored by the LaMBO model. When applied to the sequence population, the Pareto ranking method effectively selected sequences any of whose objective scores were superior to the majority of those in the initial pool (Fig. 4, Supplementary Table S2).

**Figure 4:**
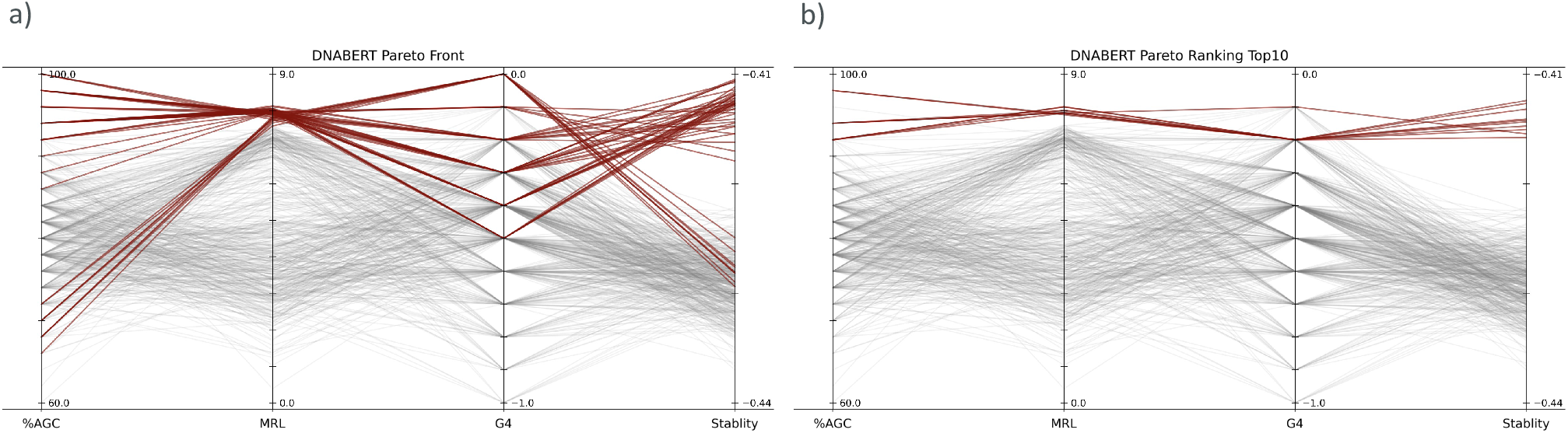
Candidate selection methods were compared using the same set of sequences after 64 rounds of LaMBO-DNABERT training. In both panels, gray lines represent all 512 sequences in the initial sequence pool (pool 3). Colored lines represent the selected candidates based on a) non-dominated solutions and b) Pareto ranking scores.

We experimentally examined 15 designed sequences, along with three 5′ UTR sequences from the initial pools, for their translation levels. Each 5′ UTR sequence was concatenated to an eGFP gene, and the *in vitro*-transcribed mRNA was transfected into 293T cells for fluorescence measurement. While the designed sequences were optimized for additional objectives, their translation levels were comparable to the top-performing sequences from the initial pools (Supplementary Figure S2). However, we unexpectedly observed a decreasing tread between the predicted MRL value and the experimentally measured fluorescence, suggesting that optimizing MRL values in the higher range may have impaired the translation level of mRNA. This observation aligns with a recent report that high ribosome loads of transfected mRNAs increased translation-dependent mRNA decay (Bicknell *et al*., 2024). Therefore, further explorations is needed to identify experimentally viable metrics to substitute the predicted MRL.

## 4 Discussion

In this study, we combined a pre-trained DNA language model, DNABERT, with LaMBO to computationally optimize human 5′ UTR sequences in a multi-objective manner. Compared to CNN and BERT, DNABERT more effectively improved the population of 5′ UTR for the four objectives: MRL, AGC content, *in vitro* stability, and G4 score. We also found that the Pareto ranking method was effective in selecting a small subset of top-performing sequences. Despite being optimized for four objectives simultaneously, the designed 5′ UTR sequences exhibited translation levels comparable to the optimal sequences in the initial pool in human cells. Our *ex vivo* experiments also indicated the need of alternative metric that quantitatively links the sequence context and translation level of 5′ UTR sequences, particularly in the higher range of predicted MRL.

In our study, the Pareto ranking method effectively selected top-performing sequences from the sequence populations. One drawback of the Pareto ranking method is that it can select sequences with similar score profile and, consequently, similar sequences. We mitigated this issue by running three independent training processes and filtering the selected sequences based on sequence identity. Two other decision-making methods, the R-method (Rao and Lakshmi, 2021) and the most-isolated Pareto solution (MIPS) score (Tamura *et al*., 2023), could also be employed. The R-method uses a rank order of objectives, provided as input, to calculate the weighted scores of solutions. This method is suitable when the relative importance of objectives is known. The MIPS score, which requires no prior assumptions, is defined by a projection free energy and favors solutions isolated from other data points, making it an automatic selection method for solutions with diverse score profiles. Both methods performed similarly on our data, although a few solutions had less optimal objective scores compared to those selected using the Pareto ranking method (Supplementary Figure S3).

Beyond 5′ UTR sequence, the entire mRNA sequence need to be optimized for biomedical applications. Although our proposed LaMBO-DNABERT pipeline has the limitation of increased computational complexity as the input length increases, the objective scores used in our study are easily applicable to the entire mRNA. By employing an alternative approach, such as combinatorial optimization of each part of the mRNA sequence, our pipeline and problem setting could lay the groundwork for the multi-objective computational design of mRNA sequences.

## Author contributions

K.Y. and M.H. designed and conceived this study and drafted the manuscript. M.H. supervised the project. K.Y. and K.S. performed computational experiments, analyzed data, and prepared the figures. N.A., K.H., S.T., M.I., F.H., and H.A. designed and conducted the biological experiments and helped with drafting the manuscript.

## Acknowledgements

We are grateful for the insightful discussions with members of the AMED/LEAP project, particularly Dr. Satoshi Uchida and Dr. Yoshihiro Shimizu. Once the manuscript was drafted, ChatGPT-4o (OpenAI) was used to check grammatical errors and improve sentence clarity. Every change suggested by ChatGPT-4o was manually reviewed by K.Y.

## Supplementary data

Supplementary methods, tables 1-2, and figures 1-3 are also available. The source code used in this study is available at https://github.com/hmdlab/lambo5utr.

