## Supplementary Materials for "Multi-objective computational optimization of human 5′ UTR sequences"

### Supplementary Methods

#### 1. Construction of IVT templates by PCR

Using plasmid DNA containing the eGFP sequence (Sample *et al.*, 2019), IVT templates were prepared through PCR using the KOD-Plus-Neo kit (Toyobo). The sequence of the templates (952 bp) is shown below. The templates included a T7 promoter (underlined), a specific 50-nt 5′ sequence ($N_{50}$), the eGFP coding sequence (bold), and a poly-$A_{70}$ tail.

5′ TTATCGAAATTAATACGACTCACTATAGGGACATCGTAGAGAGTCGTACTTA[$N_{50}$] **ATGGGCGAATTAAGT AAGGGCGAGGAGCTGTTCACCGGGGTGGTGCCCATCCTGGTCGAGCTGGACGGCGACGTAAACG GCCACAAGTTCAGCGTGTCCGGCGAGGGCGAGGGCGATGCCACCTACGGCAAGCTGACCCTGAA GTTCATCTGCACCACCGGCAAGCTGCCCGTGCCCTGGCCCACCCTCGTGACCACCCTGACCTACG GCGTGCAGTGCTTCAGCCGCTACCCCGACCACATGAAGCAGCACGACTTCTTCAAGTCCGCCATG CCCGAAGGCTACGTCCAGGAGCGCACCATCTTCTTCAAGGACGACGGCAACTACAAGACCCGCGC CGAGGTGAAGTTCGAGGGCGACACCCTGGTGAACCGCATCGAGCTGAAGGGCATCGACTTCAAG GAGGACGGCAACATCCTGGGGCACAAGCTGGAGTACAACTACAACAGCCACAACGTCTATATCAT GGCCGACAAGCAGAAGAACGGCATCAAGGTGAACTTCAAGATCCGCCACAACATCGAGGACGGC AGCGTGCAGCTCGCCGACCACTACCAGCAGAACACCCCCATCGGCGACGGCCCCGTGCTGCTGCC CGACAACCACTACCTGAGCACCCAGTCCAAGCTGAGCAAAGACCCCAACGAGAAGCGCGATCACA TGGTCCTGCTGGAGTTCGTGACCGCCGCCGGGATCACTCTCGGCATGGACGAGCTGTACAAGTTC GAATAA** AGCTAGCGCCTCGACTGTGCCTTCTAGTTGCCAGCCATCTGTTGTTTG[$A_{70}$] 3′

In the first PCR, an individual forward (Fw) primer (5′ CATCGTAGAGAGTCGTACTTA[$N_{50}$]ATGGGCGAATTAAGTA AGGGC 3′ ) and a universal reverse (Rev) primer (5′ CAAACAACAGATGGCTGGCA 3′ ) were used to amplify the sequence containing the eGFP coding region. The PCR was performed in accordance with the conditions recommended by the supplier. The thermal cycling conditions were as follows: 98 °C, 2 min → (98 °C, 10 s → 68 °C, 1 min)× 25 cycles → 68 °C, 5 min. After digestion of the template DNA with SpeedCut Dpn I (Rikaken), the PCR product (851 bp) was purified using the Wizard SV Gel and PCR Clean-Up System (Promega). The second PCR was performed on the first PCR product using a universal forward primer containing a T7 promoter (5′ TTATCGAAATTAATACGACTCACTATAGGGACATCGTAGAGAGTCGTACTTA 3′ ) and a reverse primer with a poly-dT tail (5′ [$T_{70}$]CAAACAACAGATGGCTGGCA 3′ ) using following thermal cycling protocol: 98 °C, 2 min → (98 °C, 10 s → 68 °C, 1 min)× 25 cycles → 68 °C, 5 min. All PCR reactions were analyzed by agarose gel electrophoresis, and the 5′ sequences in the second PCR product were confirmed by Sanger sequencing using the ABI PRISM 3500xL Genetic Analyzer (Center for Gene Research, Nagoya Univ.).

#### 2. Preparation of mRNAs

5′ capped mRNAs were prepared by IVT using a hydrophobic-tagged dinucleotide cap analog, DiPure (Inagaki *et al.*, 2023). The capped mRNAs were transcribed in a reaction mixture consisting of 5 ng/µL template DNA (PCR product), 2 mM NTPs, 2 mM DiPure cap analog, 40 mM Tris-HCl (pH 8.0), 8 mM $MgCl_2$, 2 mM spermidine, 5 mM DTT, 0.002 U/µL pyrophosphatase (prepared in-house) and 10 U/µL T7 RNA polymerase (prepared in-house). After incubation at 37 °C for 2 h, DNase (Takara Bio) was added to the reaction at a final concentration of 1 U/µL, followed by an additional 15 min incubation at 37 °C. The transcript was recovered by LiCl precipitation, and capped RNA was isolated using ion-paired reversed-phase high-pressure liquid chromatography (IP-RP-HPLC) with a YMC-TriartBio C18 column (YMC corp., 250 × 4.6 mm I.D.) on a Shimadzu Prominence HPLC system (pump, LC-20AD; detector, SPD-M40). Solution_A (100 mM TEAA (pH 7.0) containing 5 % acetonitrile) and Solution_B (100 mM TEAA (pH 7.0) containing 50 % acetonitrile) were used at a flow rate of 1 mL/min. The content of Solution_B was increased from 15 to 30 % over 20 min. The column temperature was maintained at 50 °C. Uncapped and capped mRNA were eluted at 11.5 and 15.0 min, respectively, and fully separated. The collected capped mRNA was irradiated with 365-nm light at 4.0 mW/$cm^2$ for 10 min to remove the hydrophobic tag (MAX-350 compact xenon light source, Asahi Spectra). The capped RNA was further purified by RP-HPLC under the same conditions described above. The purity of the 18 mRNAs was confirmed by denaturing polyacrylamide gel electrophoresis and a dot blot test using anti-dsRNA J2 antibody (Jena Bio).

#### 3. Translation activity measurement of mRNAs in cultured 293T cells

293T cells (Riken Cell Bank, RCB2202) were cultured in DMEM (Wako) supplemented with 10 % fetal bovine serum (Invitrogen) at 37 °C in a 5 % $CO_2$ atmosphere. mRNA (100 ng per well) was mixed with Lipofectamine MessengerMAX reagent (Thermo, 0.15 µL per well) in Opti-MEM I Reduced Serum Media (Thermo, 10 µL per well) in a 96-well black cell culture plate. After a

10-min incubation, 293T cells suspended in growth media without phenol red ($2.0 \times 10^5$ cells/mL, 100 µL per well) were added to the RNA/Lipofectamine mixture in the plate. eGFP expression from the mRNAs was monitored using a Spark Microplate Reader (Tecan) with $CO_2$ humidity control at 37 °C. Fluorescence (excitation at $(485 \pm 10)$ nm, emission at $(535 \pm 10)$ nm) was measured at 30-min intervals for 24 hours post-transfection. All values were summed to determine the mRNA expression level.

### Supplementary Tables and Figures

Supplementary Table S1: Tuned hyperparamaters of the models.

| Hyperparameter | LaMBO-CNN | LaMBO-BERT | LaMBO-DNABERT |
|---|---|---|---|
| Learning rate | 0.003 | 0.003 | 0.3 |
| Entropy penalty | 0.003 | 0.03 | 0.3 |
| Mask ratio | 0.3 | 0.3 | 0.2 |

Supplementary Table S2: Selected 5′ sequences and their predicted scores. Five sequences were selected as described in the method section of the manuscript. Additionally, three sequences were selected from the initial sequence sets using the same selection process.

| Index | Sequence | %AGC | MRL | G4 | Stability |
|---|---|---|---|---|---|
| CNN#1 | CACUAGGAAGAGAAGAGAAAAGAAAAUACACAAAUACAAAGGAGCAAACC | 94 | 7.895 | -0.1 | -0.417 |
| CNN#2 | CAGAGAGAAAGCACACAGGAAGACACAAGGACAAGACGCAGAGGAUCCAA | 98 | 7.654 | -0.2 | -0.422 |
| CNN#3 | CAGACAGCAGUAGAGAGACAGAUCCUAACCCAGAAACACAAACAUCUGCU | 88 | 7.804 | -0.2 | -0.421 |
| CNN#4 | CAAAAAAAGAAAAAGAACCAACAGUUGGACCCCAAAUCCUCUGCAAUACA | 88 | 7.715 | -0.1 | -0.421 |
| CNN#5 | CACAAGGAGUAGAACAAAUUCACACAAGCGAAAGAGUACACUUCACUGAU | 84 | 7.946 | -0.2 | -0.418 |
| BERT#1 | CCGAGAACAAAAACGAAAGAGAGAGAGAACACUGAGCUAAGCUGAGCAAC | 94 | 7.763 | -0.1 | -0.421 |
| BERT#2 | ACAAAAGGACAAAACAGAACAGAGAACAGCAGUCGCAAACAGUAGAUACC | 94 | 7.764 | -0.2 | -0.419 |
| BERT#3 | CUUAAAGGAACAAACGAAAAAGGAAAGGAAGCAAAUCCAUGACCAAGGAC | 92 | 7.761 | -0.2 | -0.421 |
| BERT#4 | CUUAAAACAGCAUCAAAGAGAGAAAAAGAAACGUAACAAACCAUAGAGAC | 90 | 7.678 | -0.1 | -0.418 |
| BERT#5 | ACCGGAGCCACAAACGACAGAGGAAAAAGAAGGAGGCACACAUAGACAAC | 98 | 7.605 | -0.2 | -0.422 |
| DNABERT#1 | AAGAGGAAAAAAAAAAAACAACGAAGAAAAAAAAAGAGAGAAAGAGAAUAAA | 98 | 7.906 | -0.2 | -0.413 |
| DNABERT#2 | AAGAAAGAAAAAAAAAAACAAAGAAGAAAACAAAGAGACAAAGAAAAAAAA | 100 | 7.819 | -0.2 | -0.411 |
| DNABERT#3 | GAAAAACACAGAGAAACAAUAGUAGAGGAGCCACAUACAGAACACAAGAG | 94 | 7.911 | -0.1 | -0.420 |
| DNABERT#4 | AAGAGAAACACAAAACAGAGAAACACAAGACAAAGACGAACAGAAGUAAA | 98 | 7.887 | -0.2 | -0.418 |
| DNABERT#5 | AAAGAGACAAAGAGAGAAAAGGGACCAAGAAAACUAAGACGAACAAAACC | 98 | 7.766 | -0.1 | -0.418 |
| Init#1 | AGUUCACCAAAGAAGGAAAUUCACAGCUGCAAAUGAGCACGGCAAACAAA | 88 | 7.441 | -0.2 | -0.426 |
| Init#2 | CCCAAAGCCAAUACGAAGCAACAGUUGGAUCCUAAGUCCUCUGAAAAACA | 84 | 7.581 | -0.3 | -0.423 |
| Init#3 | CCCACCAUUUCGACCAAACGAACAUCAACCAAACAAGACACUGUGUAACA | 86 | 7.159 | -0.3 | -0.425 |

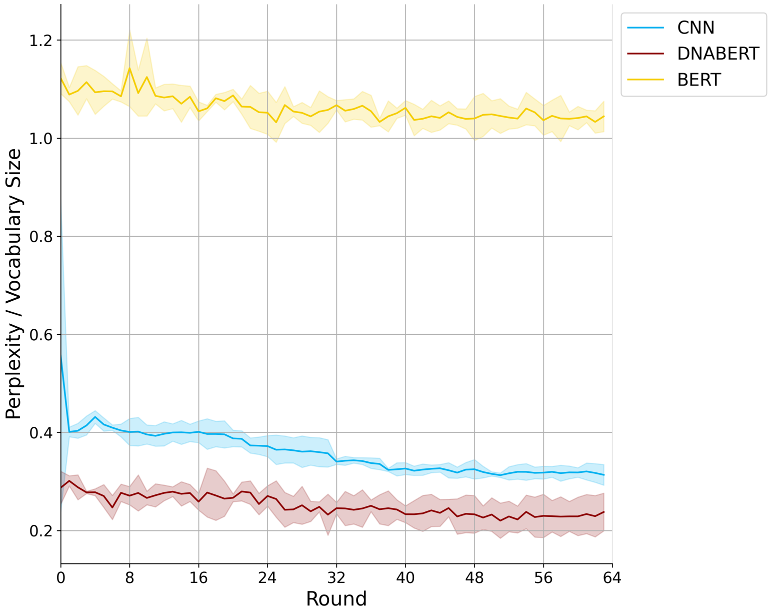

Supplementary Figure S1: Normalized perplexity of each model during training. Perplexity was divided by the vocabulary size of each model, which was 10 for CNN and 69 for BERT and DNABERT.

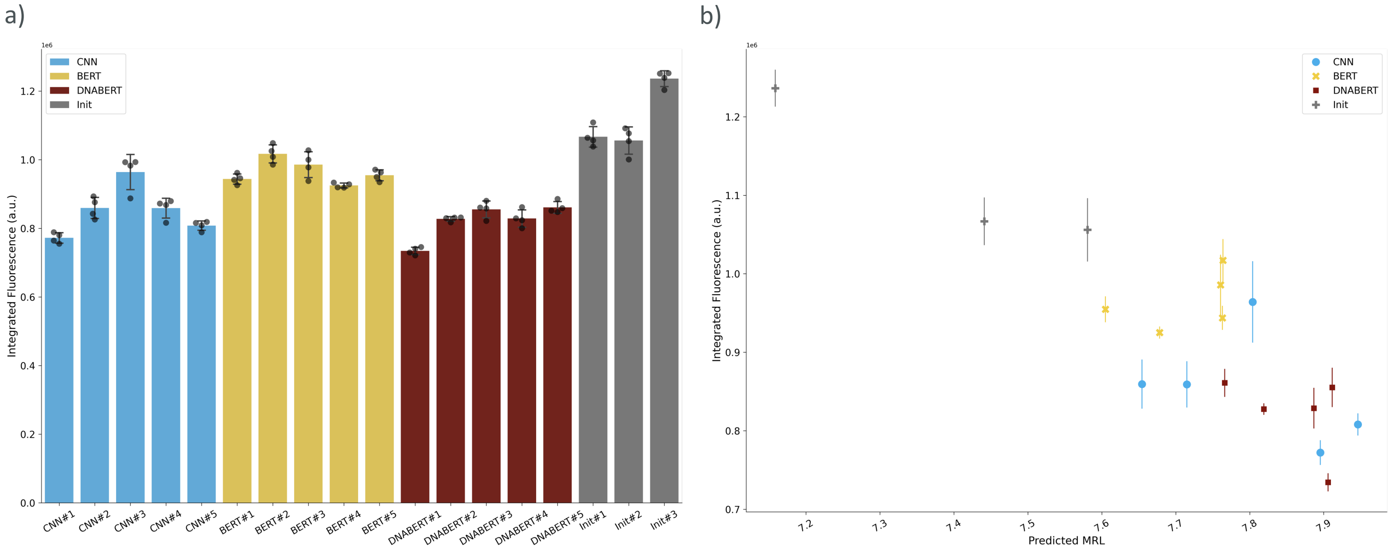

Supplementary Figure S2: The designed 5′ sequences were concatenated with an eGFP coding sequence, transfected into HEK293T cells, and GFP fluorescence was measured to evaluate translation levels. a) Fluorescence was measured over a 24-hour period post-transfection and summed over time. Each dot represents a single data point, and error bars indicate the mean and standard deviation across four replicates. The x-axis labels correspond to those in Supplementary Table S2. b) Relationship between the predicted MRL by Optimus 5-Prime and the measured fluorescence. Each marker represents the mean value, with error bars indicating the mean and standard deviation across four replicates as in a).

Comparison of decision-making methods, including Pareto optimal solutions, Pareto-ranking method, R-method, and MIPS scoring. Gray lines in both panels represent all 512 sequences in the initial sequence pool (pool 3). Colored lines represent the selected candidates based on each decision-making method.

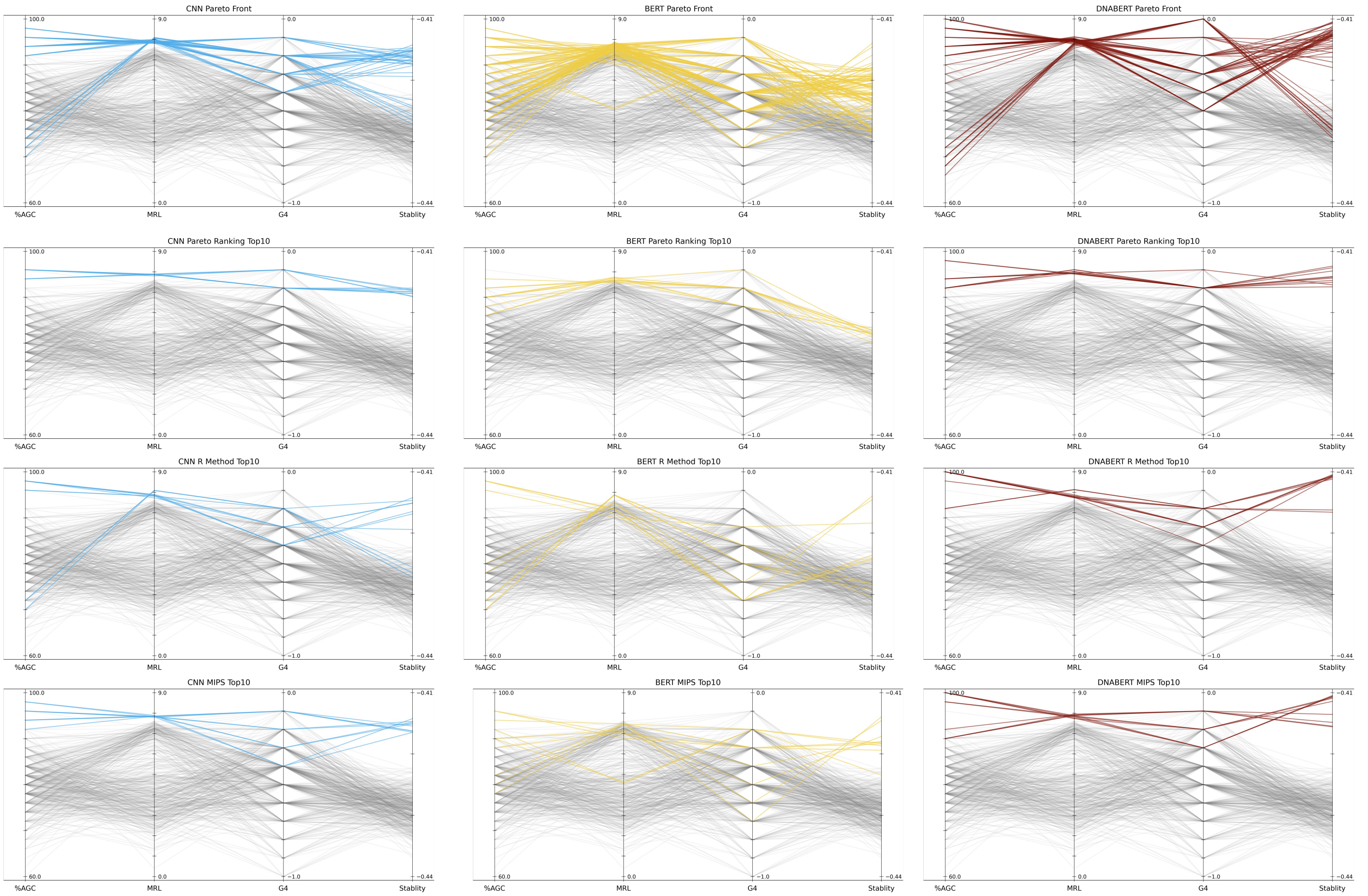

Supplementary Figure S3: Comparison of decision-making methods, including Pareto optimal solutions, Pareto-ranking method, R-method, and MIPS scoring. Gray lines in both panels represent all 512 sequences in the initial sequence pool (pool 3). Colored lines represent the selected candidates based on each decision-making method.
